# Forced proximity promotes the formation of enduring cooperative relationships in vampire bats

**DOI:** 10.1101/2022.01.30.478382

**Authors:** Imran Razik, Bridget K. G. Brown, Gerald G. Carter

**Affiliations:** The Ohio State University, Columbus, Ohio, United States; Smithsonian Tropical Research Institute, Balboa Ancón, Panamá, Panama

**Keywords:** vampire bats, cooperation, social relationships, cooperative relationships, association

## Abstract

Spatial assortment can be both a cause and consequence of cooperation. Proximity promotes cooperation when individuals preferentially help nearby partners, and conversely, cooperation drives proximity when individuals move towards more cooperative partners. However, these two causal directions are difficult to distinguish with observational data. Here, we experimentally test if forcing randomly selected pairs of equally familiar female common vampire bats (*Desmodus rotundus*) into close spatial proximity promotes the formation of enduring cooperative relationships. Over 114 days, we sampled 682 hours of interactions among 21 females captured from three distant sites to track daily allogrooming rates over time. We compared these rates before, during, and after a one-week period, during which we caged random triads of previously unfamiliar and unrelated vampire bats in close proximity. After the week of close proximity when all bats could again freely associate, the allogrooming rates of pairs forced into proximity increased more than those of the 126 control pairs. This work is the first to experimentally demonstrate the causal effect of repeated interactions on cooperative investments in vampire bats. Future work should determine the relative importance of mere association versus interactions (e.g. reciprocal allogrooming) in shaping social preferences.

## Introduction

Spatial assortment can both be a cause and a consequence of cooperation. This principle applies to both ecological and evolutionary timescales [1–4], and across a great diversity of social life from microbes [5] to humans [6]. Proximity drives cooperation when individuals primarily help nearby partners, and conversely, cooperation drives proximity when individuals move towards more cooperative partners. Although both forces are predicted by theory, these two causal directions are not easily distinguished with observational data.

Individuals forced to associate might form enduring cooperative relationships that persist beyond the period of forced proximity, similar to how randomly assigned first-year college roommates are more likely to be friends after graduating [7]. Several past results suggest that such a phenomenon occurs in common vampire bats (*Desmodus rotundus),* a species where preferred associations correlate with cooperative interactions, such as allogrooming and regurgitated food sharing [8–12]. Vampire bats housed together for months in captivity appeared to develop stronger social preferences for each other and then maintained these associations for at least eight days after being released back into the wild [11]. Female pairs that were introduced to one another in small cages started food-sharing relationships faster than pairs that first met in a larger cage with more options for social partners [12]. In field sites where vampire bats switch among roosts, pairwise roost-sharing rates predict allogrooming and food sharing [8,9,13]. However, all these studies lack the proper experimental controls to show a causation.

Here, we experimentally tested if forcing proximity among randomly selected triads of female common vampire bats (*Desmodus rotundus*) promotes the formation of enduring cooperative relationships. We tracked changes in allogrooming because it is a cooperative and highly symmetrical investment of time and energy that is (*i*) directed to individuals in need [14], (*ii*) sufficiently frequent during the formation of new relationships [12], and (*iii*) used by individuals to establish and maintain more costly food-sharing relationships [12]. We found that just one week of forced proximity between recently introduced and unrelated females increased their allogrooming rates, relative to a control group, after the manipulation ended.

## Materials and Methods

### Study colony

Experimental subjects were 21 individually marked female common vampire bats in a captive colony. Seven bats were sourced from each of three different sites in Panama (Lake Bayano, Tolé, or La Chorrera) that were 120-350 km apart, and housed in an outdoor flight cage at the Smithsonian Tropical Research Institute in Gamboa, Panamá. See Razik et al. [15] and the supplementary information for more details on the captive colony.

### Experimental design

Subjects experienced three phases: pre-treatment (six weeks), forced proximity (one week), and post-treatment (nine weeks). During the pre-treatment phase, all bats could freely interact in a flight cage (2.1 × 1.7 × 2.3 m). During the treatment phase, we randomly assigned female bats into seven triads (including one bat from each site), each housed in a 28 x 28 x 40 cm clear, acrylic observation cage. For the post-treatment phase, we released all bats back into the flight cage and monitored interactions (Figure 1). We classified the 21 recently introduced pairs that were forced into proximity as ‘test dyads’, the other 126 recently introduced pairs as ‘control dyads’, and the 63 pairs of bats caught from the same site as ‘familiar dyads’.

**Figure 1.**
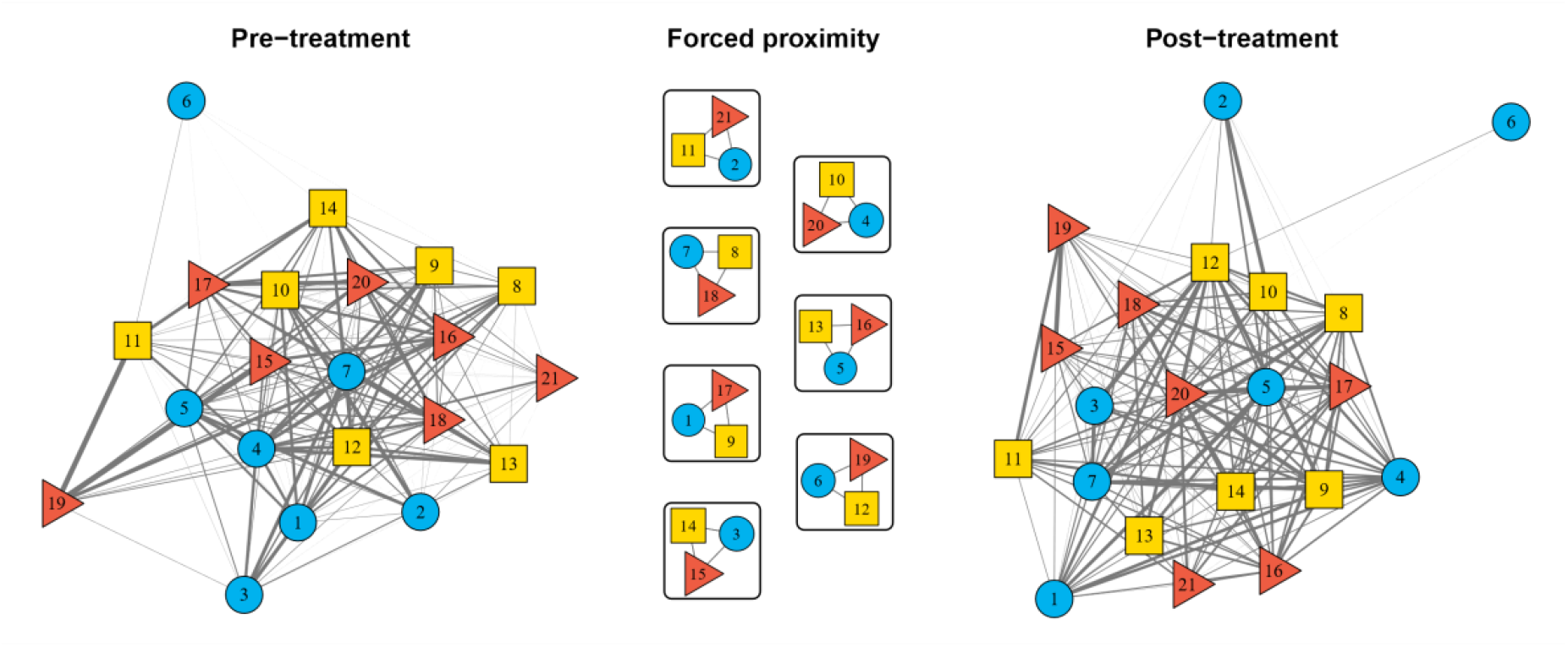
Experimental design. Nodes are bats.Node color/shape shows capture site (Lake Bayano = blue circles, Tolé = yellow squares, La Chorrera = red triangles). Width of grey edges shows the relative allogrooming log rates during the six-week pre-treatment phase, the one-week of forced proximity, and the nine-week post-treatment phase.

To measure allogrooming, we used three infrared surveillance cameras (Foscam NVR Security System) to sample all social interactions among bats for three to six hours each day from 23 June 2019 to 14 October 2019. Over 114 days, we sampled 682 hours of interactions, recording allogrooming bouts that were at least five seconds in duration, and noting both the actor and receiver. To measure dyadic allogrooming rates, we averaged the seconds of allogrooming per hour across all hours in which the actor and receiver could interact, first averaging the rates in both directions (most allogrooming is bidirectional), then transforming these values with *natural log (x + 1),* because the variance of allogrooming duration increases with the mean. We call these measures “allogrooming log rates”.

### Statistical analyses

To test for an effect of forced proximity, we compared the mean change in allogrooming log rates from the pre-treatment to the post-treatment phase for test dyads, control dyads, and familiar dyads, then calculated 95% confidence intervals (CI) for all mean changes using bootstrapping (percentile method in the *boot* R package) [16]. Using the same bootstrapping method, we calculated the 95% CI for the mean change in the proportion of allogrooming log rates that bats directed to familiar versus previously unfamiliar partners.

To test if the mean change in allogrooming log rates differed between test and control groups more than expected by chance, we used a permutation test. We compared the observed difference to a distribution of 5000 expected differences obtained from running the same analysis after randomly re-assigning bats from different capture sites into possible new forced-proximity triads. To check the robustness of our original result from this analysis, we also conducted several alternative analyses (see supplementary information). Briefly, our results hold when (*i*) excluding 17 control dyads with pre-treatment allogrooming rates higher than the maximum rate observed in test dyads (to account for regression to the mean effects), (*ii*) removing two bats that were sampled fewer times than other females, (*iii*) removing bats that had a Staph infection during the pre-treatment period, and (*iv*) drawing inferences from a mixed-effect model. All analyses confirm the original result of a clear difference between test and control groups.

To test if allogrooming log rates in the 21 test dyads during the forced-proximity phase predicted changes in allogrooming log rates between the pre-treatment and post-treatment phase, we fit a linear model with the change in allogrooming as a ranked response and forced-proximity allogrooming ranks as the effect. To test if forced-proximity allogrooming ranks predicted allogrooming during the post-treatment phase after controlling for baseline levels, we fit a linear model with post-treatment allogrooming rank as a response and both pre-treatment and forced-proximity allogrooming ranks as effects. Alongside parametric p-values, we present one-sided p-values from a permutation test (‘permutation-p’) comparing the observed coefficients to those expected when allogrooming rates are randomized within each forced-proximity cage (5000 permutations).

## Results

From the pre-treatment to the post-treatment phase, bats increased the proportion of their allogrooming directed to bats from different capture sites (mean increase = 0.06, 95% CI = [0.002, 0.11]. Allogrooming increased more in the test dyads that were forced into proximity compared to the control dyads (Figure 2; mean increase for test dyads = 0.66 log s h^−1^ [0.37, 0.97], for control dyads = 0.17 log s h^−1^ [0.04, 0.30]; difference = 0.50 log s h^−1^, p = 0.002). This effect was evident throughout the post-treatment period (Figure S1, supplementary information).

**Figure 2.**
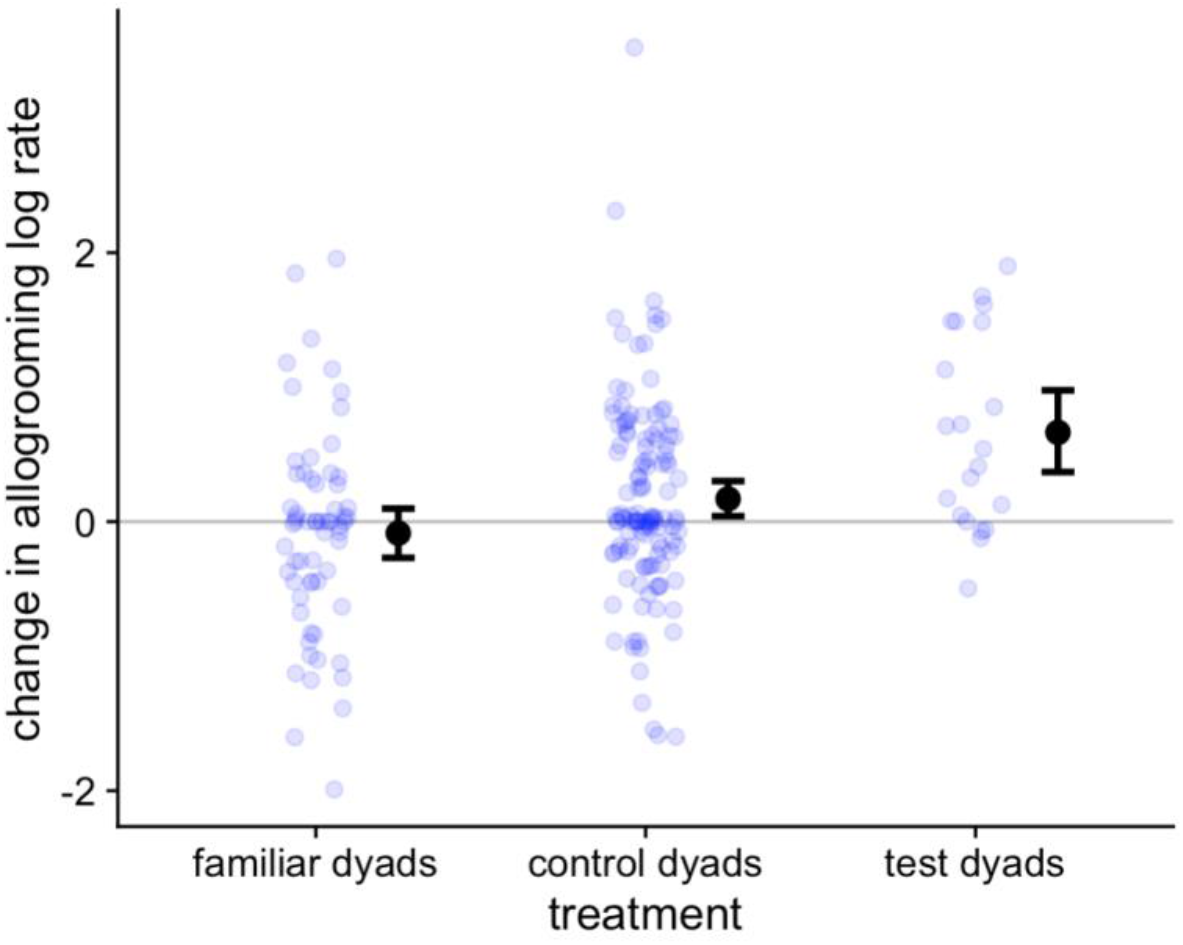
Changes in allogrooming in familiar, control, and test dyads. Blue points are bat pairs. Larger black points are means with 95% CIs. Familiar dyads are from the same capture site. Test and control dyads are pairs of bats from different capture sites that were either forced into proximity or not, respectively.

Within the 21 test dyads, allogrooming rates during forced proximity were more clearly correlated with post-treatment rates (Spearman’s ρ = 0.49, ρ = 0.025) than pre-treatment rates (ρ = 0.33, p = 0.15); however, we failed to find clear evidence that allogrooming ranks among the 21 test dyads during the forced proximity phase predicted changes in allogrooming (β = 0.28, p = 0.206, permutation test p = 0.15), or post-treatment allogrooming when controlling for the baseline pre-treatment allogrooming (β = 0.32, p = 0.096, permutation test p = 0.074; Figure S5, supplementary information).

## Discussion

We introduced previously unfamiliar and unrelated vampire bats in captivity, allowing them to freely associate for six weeks in an outdoor flight cage, then forced randomly selected triads into proximity for one week. Over the next nine weeks, while bats could again freely associate, we found that allogrooming rates of pairs forced into proximity had increased more than those of pairs that were not forced into proximity. This finding shows that manipulating proximity can promote the formation of enduring cooperative relationships that persist beyond the manipulation period.

Animals capable of individual social recognition are likely to have a familiarity bias, leading to social preferences among individuals that happen to meet earlier in time or be closer in space. Early-life associations can be a cue for kin discrimination [e.g. 17] and lead to strong preferences later in life [18–21]. For example, newborn pipistrelle bats (*Pipistrellus kuhlii*) raised in separate groups for six weeks preferred to associate with and groom familiar conspecifics and heterospecifics after being released into a common flight cage [20,21]. Preferred associations can also develop quickly during adulthood. For instance, guppies (*Poecilia reticulata*) develop new schooling preferences for familiar individuals over just 12 days, and the effect of familiarity appears stronger than phenotypic similarity [22,23].

The mechanism underlying the change we observed remains unclear. The observed increases in allogrooming were likely caused by some unknown combination of elevated *associations* and more frequent or intense *interactions* (e.g. bats increased allogrooming with partners that groomed them more). Although we failed to detect a clear correlation between allogrooming intensity within the forced proximity period and changes in allogrooming afterwards, our power to detect this effect is far worse than our ability to detect the effect of forced proximity, because we did not manipulate grooming directly, the forced-proximity grooming rates were based on far fewer observations (one week), and we had far fewer dyads in the analysis (21 vs 147 dyads).

Furthermore, past work on vampire bats strongly suggests that spatial proximity alone cannot entirely explain either (*i*) the large variation in cooperative behavior across individuals and pairs living in close proximity in captivity [12,24] or (*ii*) the changes in cooperative relationships owing to manipulations of experience [12]. For example, allogrooming was a better predictor of new food-sharing relationships within periods of forced proximity than within periods of free association [12]—the opposite of what is expected if food sharing is caused only by forced proximity. Decisions to help are likely determined, at least in part, by partner experiences or traits. If individuals use frequent association as an honest cue of partner availability, and if availability is desirable in a social partner, then this could result in a positive feedback loop during the social bonding process, where bats preferentially groom frequent associates and preferentially associate with frequent groomers. Future experiments should disentangle the roles of interactions versus association in social bonding.

Many aspects of the social bonding process remain mysterious. If bats that were forced to associate in the same small cage similarly perceived these shared experiences as negative, would this weaken or strengthen the process of social bonding? Animals might prefer partners with whom they share rewarding experiences because of simple associative learning; alternatively, negative or stressful experiences that are shared might facilitate or reinforce partner choice [25]. For instance, guppies that were exposed to an environment where they perceived high-predation risk were found to exhibit stronger partner preferences for one another than those that were not [26]. Furthermore, to what extent is the process of social bonding influenced by the quantity and quality of other available partners? Vampire bats form new relationships faster in the absence of familiar partners [12], and the benefits of having fewer stronger bonds versus many weak ties could vary with many aspects of the social environment, such as social stability [24] and the number of alternative relationships. If social bonds require long-term cooperative investments, then do individuals have ‘investment strategies’ that depend on the marginal return on each additional unit of investment? Answering such questions will require controlling association while carefully manipulating interactions within pairs.

## Ethics

This work was approved by the Smithsonian Tropical Research Institute Animal Care and Use Committee (#2015-0501-2022) and the Panamanian Ministry of the Environment (#SEX/A-67-2019).

## Data accessibility

Data and R code are available on Figshare (https://doi.org/10.6084/m9.figshare.19077431.v2).

## Authors’ contributions

IR performed the experiments, collected the data, and helped with experimental design, project coordination, data analysis, and writing. BKGB helped with experiments and data collection. GGC conceived of the study and helped with experimental design, project coordination, resources, funding, data analysis, and writing. All authors gave final approval for publication and agree to be held accountable for the work performed therein.

## Acknowledgments

The authors thank Rachel Page for lab space and resources, the staff at the Smithsonian Tropical Research Institute, and the following individuals who assisted with vampire bat care and/or data collection: D. Aparicio, G. Cohen, L. Dück, D. Girbino, E. Kline, C. Marroquin, S. Ripperger, and S. Stockmaier.

## Supplementary information

### Supplemental details on methods

We fed bats with refrigerated or thawed cattle or pig blood that was defibrinated with 44 g of sodium citrate and 16 g of citric acid per 19 L container. Female bats that were estimated to be at least 10 months old (Wilkinson 1985; Greenhall & Schmidt 1988) were classified as “adults” while younger bats were classified as “juveniles”. From a cave at Lake Bayano, Panamá, we caught six adult females, two juvenile males, and one juvenile female. From outside a large, hollow tree roost in Tolé, Panamá, we captured seven adult females. From outside another hollow tree roost in La Chorrera, Panamá, we captured eight adult females. These sites were 120-350 km apart such that bats from different sites were safely assumed to be unrelated and unfamiliar. We identified bats using unique sets of forearm bands (Porzana and National Tag). On 14 June 2019, we released all bats into an outdoor flight cage (2.1 m x 1.7 m x 2.3 m).

On 5 July, we added two more adult females from the capture site in Tolé, Panamá. These two bats were subjects in a past experiment where they were captured in the wild and studied in captivity from December 2015 to September 2017 before being released back into the wild. The two re-captured bats were excluded as subjects during the treatment phase of our experiment (described below) due to their exposure to this previous experiment.

We sampled behavioral interactions in one-hour periods at 0400, 0500, 0900, 1900, 2000, and 2100 h, and on a few occasions, at 0100, 0300, 1100, and 2200 h. To induce and measure food sharing, we repeatedly fasted the bats and scored mouth-licking; however, we found that mouth-licking was not well correlated with mass gain, and we lacked the number of new food-sharing interactions necessary to reliably measure changes in relationships through food sharing. Therefore, we focused our analyses on allogrooming.

### Supplemental analyses

#### Results were robust when controlling for regression to the mean effects by removing high-grooming dyads from the control group

Despite randomly selecting dyads for the test group, we noticed that the sample mean (but not the median) allogrooming rate for test dyads was lower than those for the control group during the pre-treatment phase because the 17 highest grooming pairs ended up in the control group (Figure S1). The probability for a mean difference of this size or larger to occur by chance was 16% (estimated by randomly re-assigning the proximity treatment to introduced dyads 5000 times; observed mean difference = 0.67 log s h^−1^; 95 % of expected values were between 0.51 and 1.25 log s h^−1^, one-tailed p = 0.16). One issue is that this starting difference between the test and control group could lead to a larger increase in the test group simply due to regression to the mean. To control for this, we repeated our main statistical analyses after first excluding 17 control dyads wherein grooming rates during the pre-treatment phase were outside the range of grooming rates that were observed in the test dyads. After excluding these dyads from the control group, our conclusions did not change: allogrooming log rates still increased more in test dyads that were forced into proximity compared to control dyads (mean increase: test dyads = 0.66 log s h^−1^, 95% CI = [0.37, 0.97 log s h^−1^]; control dyads = 0.28 log s h^−1^, [0.16, 0.41 log s h^−1^], observed difference = 0.38 log s h^−1^, p = 0.006; Figure S1, S2).

**Figure S1.**
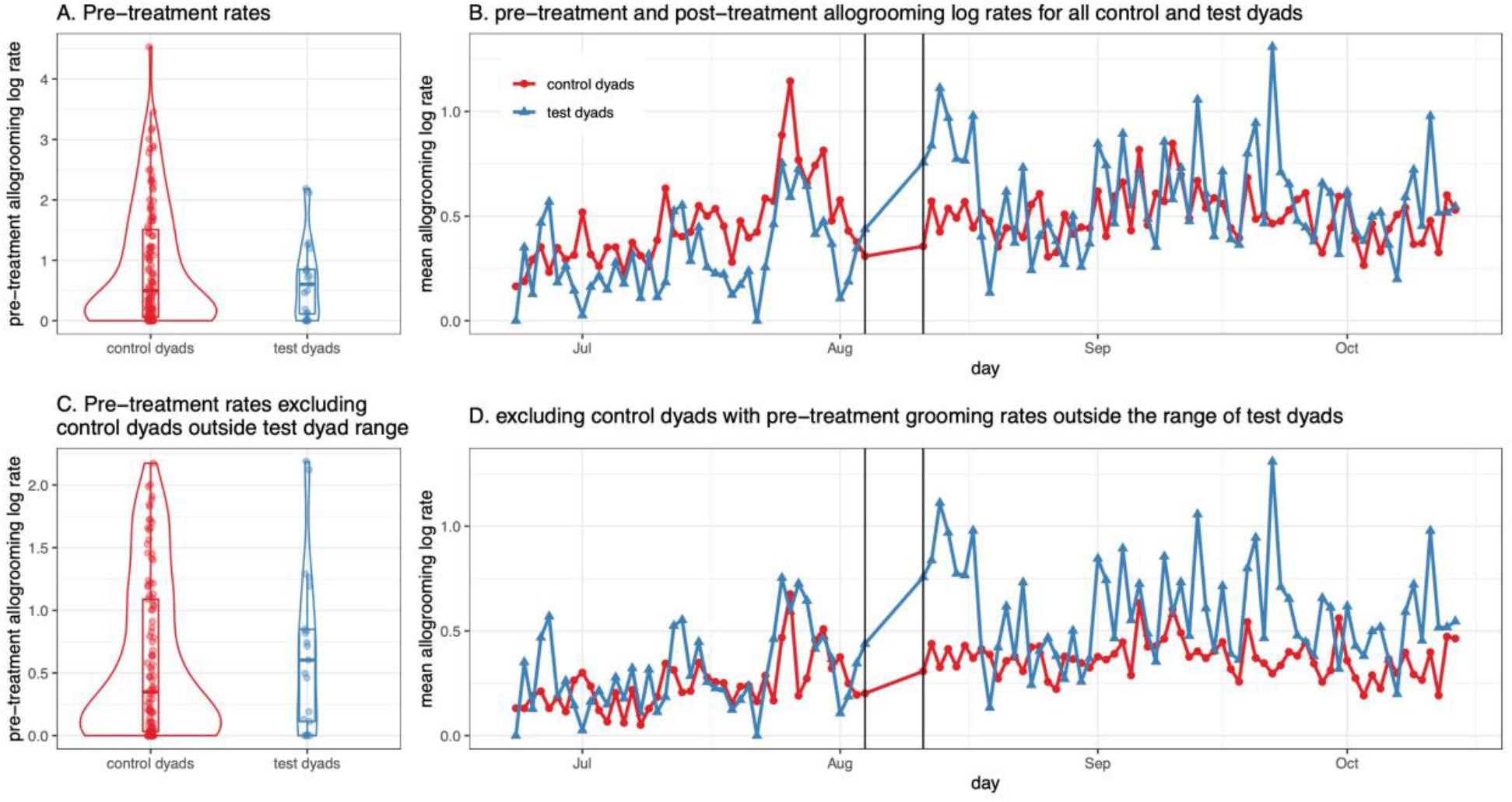
Pre-treatment and post-treatment changes in allogrooming for test and control dyads. Panel A shows the distribution of pre-treatment allogrooming log rates for the 126 control dyads and 21 randomly selected test dyads. Panel B shows the daily allogrooming rates of test and control dyads before and after the period of forced proximity shown as two black vertical lines. Panels C and D show the same plots after excluding control dyads with pre-treatment grooming rates outside the range of the test dyads to show that the greater increase in test dyad rates was not simply due to regression to the mean.

**Figure S2.**
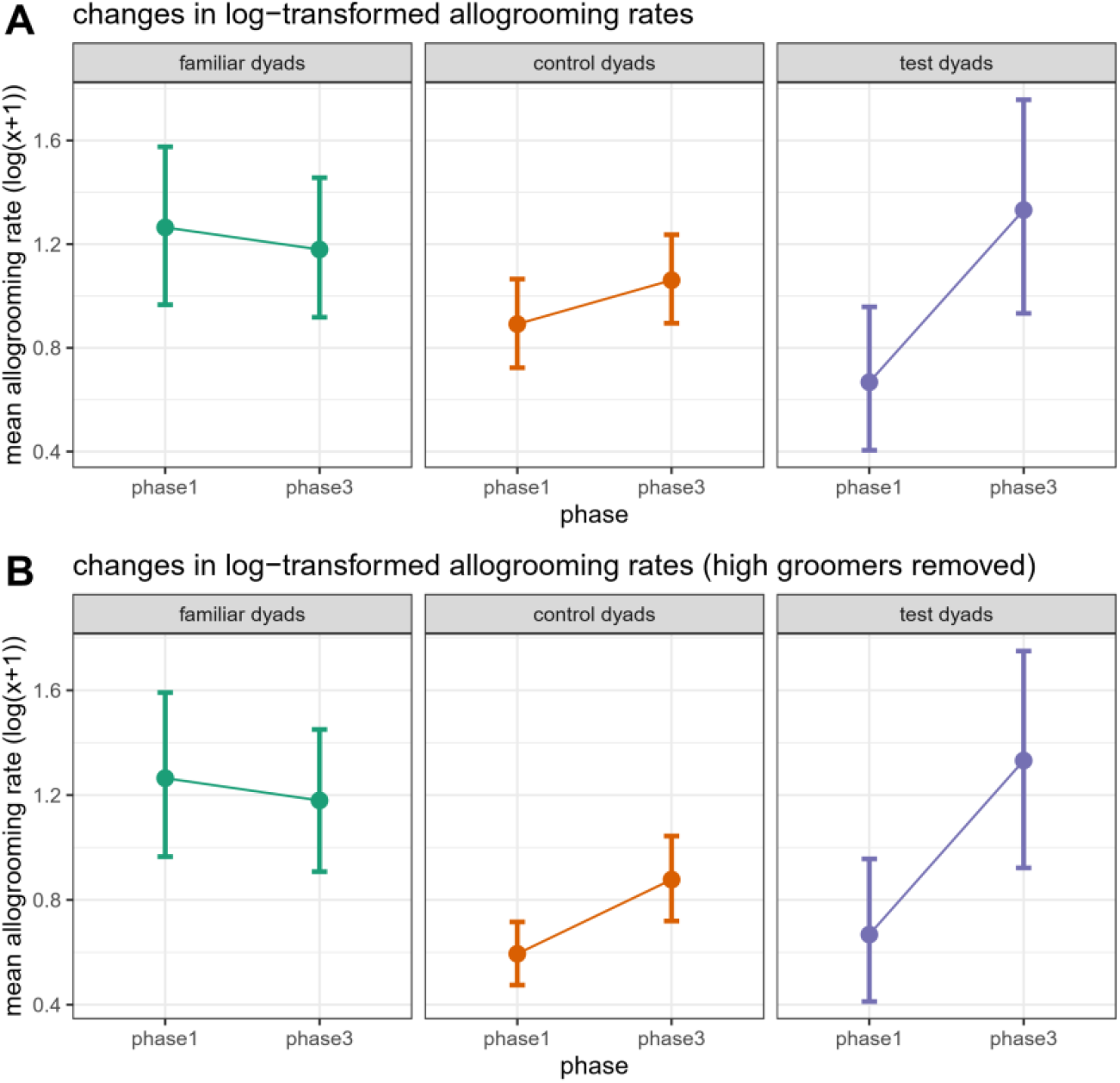
Mean grooming rates and bootstrapped 95% confidence intervals during the pre-treatment and post-treatment phase for all three dyad types. Panel A shows the main analysis comparing the change in allogrooming between dyad types. Panel B shows the alternate analysis excluding the 17 highest grooming pairs from the control group.

#### Results were robust to removing imprecise estimates for two bats

Two adult female bats that were observed during all three phases of the experiment were undersampled relative to other bats during either the pre-treatment or post-treatment phase. One bat entered the colony a month after most others, while the second bat died two weeks after the treatment phase. Given that the mean grooming rates for these bats are less precise than our grooming rates for other bats, we chose to repeat our analyses without these two bats. Our finding did not change: allogrooming log rates still increased more in test dyads that were forced into proximity compared to control dyads (mean increase: test dyads = 0.72 log s h^−1^, 95% CI = [0.40, 1.05 log s h^−1^]; control dyads = 0.21 log s h^−1^, [0.07, 0.36 log s h^−1^], observed difference = 0.52 log s h^−1^, p = 0.003).

#### Results were robust to removing estimates for bats that experienced a bacterial infection

During the pre-treatment phase of the study, nine bats developed a *Staphylococcus* infection. We therefore isolated these bats from all other bats in the colony and treated them with enrofloxacin antibiotic between 21 July and 5 August 2019. The bats recovered after administering antibiotic, but four bats that were severely infected had permanent tissue damage in their wrists and could not maintain powered, sustained flight throughout the remainder of the study. Given that these events could have affected the frequency or duration of social interactions among bats in the colony, we chose to repeat our analyses after first excluding either (1) all bats that received antibiotic treatment, or (2) the four bats that were physically impaired because of their infection.

Our results did not change when the excluding nine bats treated with antibiotic: allogrooming log rates still increased more in test dyads that were forced into proximity compared to control dyads (mean increase: test dyads = 0.65 log s h^−1^, 95% CI = [0.12, 1.18 log s h^−1^]; control dyads = −0.03 log s h^−1^, [-0.21, 0.14 log s h^−1^], observed difference = 0.68 log s h^−1^, p = 0.004). Our results also did not change when only excluding four bats that were physically impaired from their infection: allogrooming log rates still increased more in test dyads that were forced into proximity compared to control dyads (mean increase: test dyads = 0.57 log s h^−1^, 95% CI = [0.26, 0.94 log s h^−1^]; control dyads = –0.05 log s h^−1^, [– 0.19, 0.08 log s h^−1^], observed difference = 0.62 log s h^−1^, p = 0.001).

#### Alternative analysis using a mixed effect model

We also tested for an effect of forced proximity on allogrooming rates by fitting a general linear mixed-effects model. Standardized (z-transformed) post-treatment allogrooming log rates were the response. Fixed effects were dyad type (i.e., control or test dyad) and standardized pre-treatment allogrooming log rates. Random intercepts were actor and receiver identity. Although the distribution of model residuals was not normal (Shapiro-Wilk test: W = 0.98, p = 0.003), residuals were not dramatically skewed or non-normal. Consistent with our original analysis, the model suggested that allogrooming log rates were higher among test dyads than control dyads during the post-treatment phase (β =0.42, p < 0.0001), even when controlling for baseline pre-treatment levels (β = 0.61, p < 0.0001, Figure S3). Note that this analysis makes more assumptions about error distributions and independence than our permutation-based null model which simulates random assignments of bats into possible forced-proximity triads.

**Figure S3.**
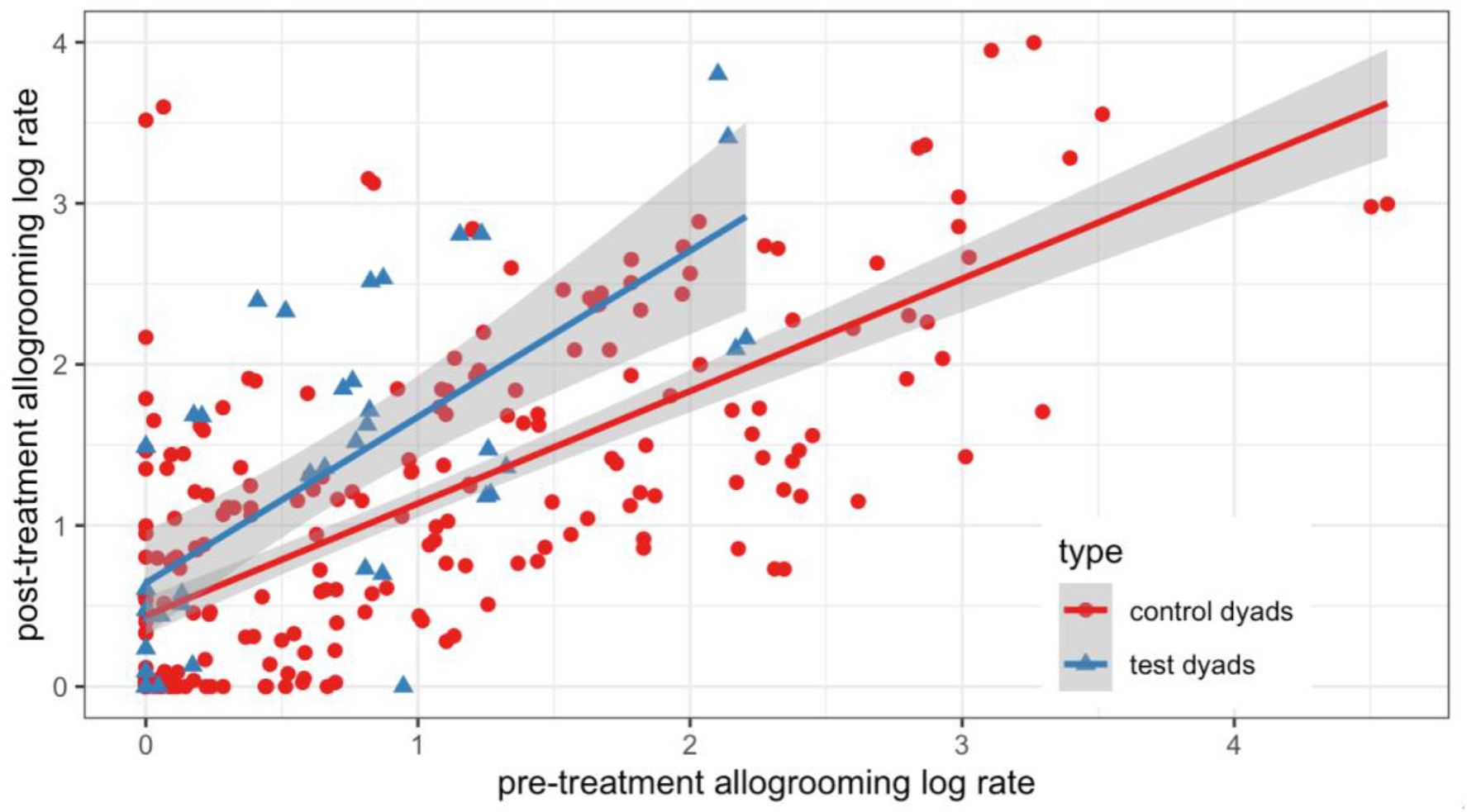
Test dyads had higher post-treatment allogrooming log rates than control dyads when controlling for baseline pre-treatment allogrooming log rates.

#### Mouth-licking rates were too low and imprecise to track cooperative relationship changes over time

To induce food sharing during the pre-treatment and post-treatment phases, we individually isolated up to three bats each day (one bat per capture site) and fasted them for 23 hours. Each bat was fasted 3-15 times, and bats were primarily fasted once every 7 days. Bats that were fasted on the same day were unable to share food with one another during a fasting trial. During the treatment phase, we fasted each bat up to an additional two times, with seven bats being fasted on a single day (one bat per small cage). When we repeated our analysis using mouth-licking rather than allogrooming rates, we saw a sample mean increase in the same direction as allogrooming, but the mean estimates were highly variable and imprecise due in part to the highly zero-inflated data (Figure S4). There are several important caveats to interpreting this analysis. First, much of mouth-licking occurred outside fasting trials and might not represent food sharing. Unlike previous studies (Carter & Wilkinson 2013, 2015; Carter et al. 2020), mouth-licking rates for fasted bats in this study were only weakly correlated with mass gain (Spearman’s ρ = 0.32, p = 0.01), possibly because some of the mouth-licking was unsuccessful begging rather than food sharing, and because fasted bats were not followed with a handheld camera as in the past study, so more food sharing likely occurred ‘offscreen’. Furthermore, only 8% of introduced dyads showed any mouth-licking during fasting trials, compared to 57% that did show allogrooming over the 4 months. In a previous study that was 14 months long, only 16% of introduced adult female dyads showed mouth-licking during fasting trials (Carter et al. 2020). For all these reasons, allogrooming rates are much better than mouth-licking rates as a measure for the development of new cooperative relationships.

**Figure S4.**
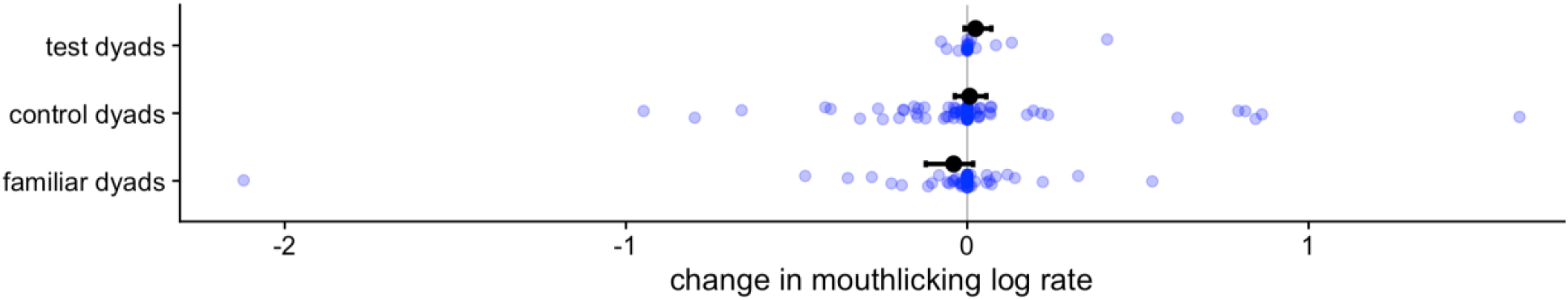
Mean change in mouth-licking rates and bootstrapped 95% confidence intervals during the pre-treatment and post-treatment phase for all three dyad types.

**Figure S5.**
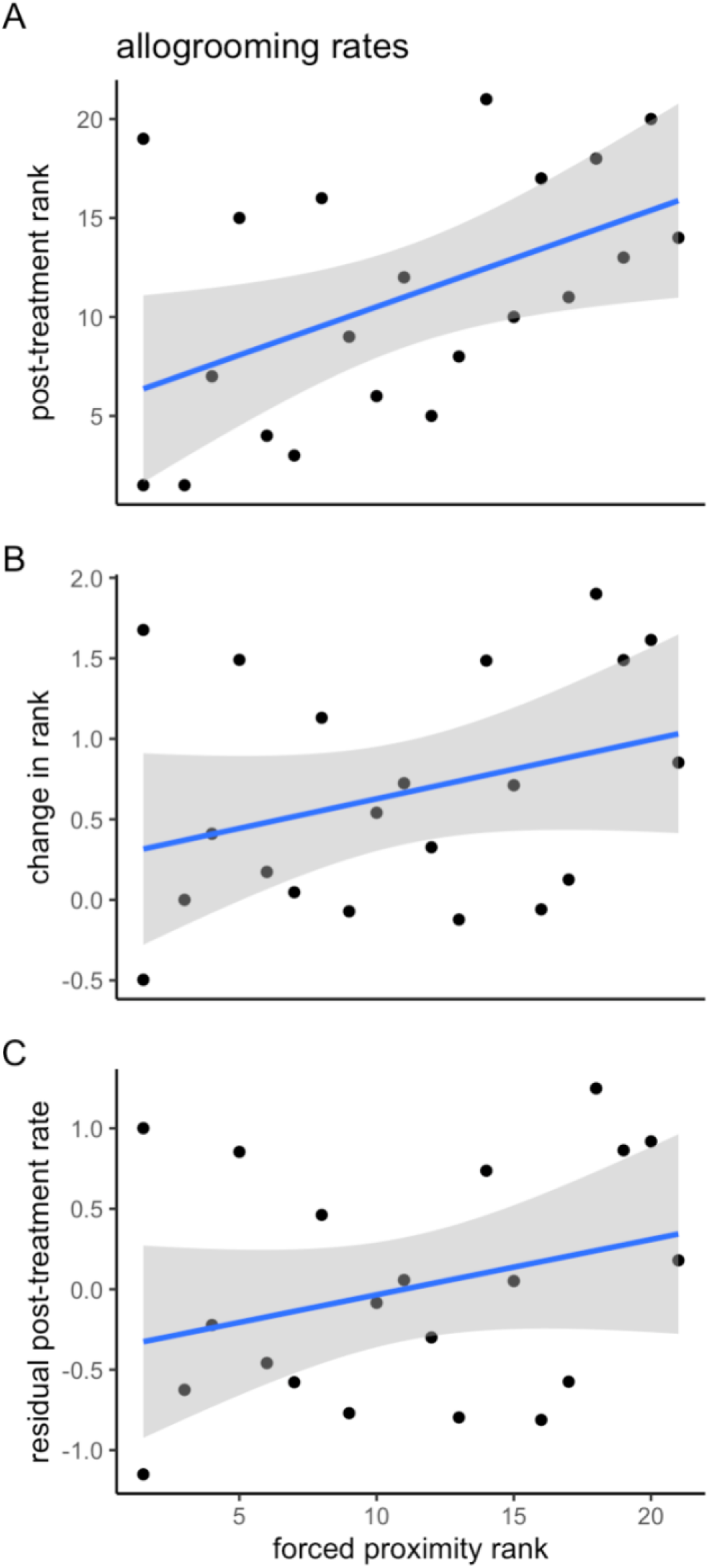
No clear evidence for effects of variation in forced-proximity allogrooming rates on post-treatment allogrooming within the 21 test dyads. Plots show linear fits (blue line) and confidence intervals on slopes (grey) between ranks of dyadic allogrooming during forced proximity and either (A) post-treatment allogrooming ranks, (B) the change in dyadic allogrooming rank from pre-treatment to post-treatment, or (C) the residuals from a linear model with pre-treatment log rates predicting post-treatment log rates (R^2^ = 0.485, p = 0.0005).

